# Genomic insights into adaptive divergence and speciation among malaria vectors of the *Anopheles nili* group

**DOI:** 10.1101/068239

**Authors:** Caroline Fouet, Colince Kamdem, Stephanie Gamez, Bradley J. White

**Affiliations:** Department of Entomology, University of California, Riverside, CA 92521; Center for Disease Vector Research, Institute for Integrative Genome Biology, University of California, Riverside, CA 92521

**Keywords:** *Anopheles nili*, divergent selection, high-*F*_ST_ regions, speciation

## Abstract

Ongoing speciation in most African malaria vectors gives rise to cryptic populations, which differ remarkably in their behaviour, ecology and capacity to vector malaria parasites. Understanding the population structure and the drivers of genetic differentiation among mosquitoes is critical for effective disease control because heterogeneity within species contribute to variability in malaria cases and allow fractions of vector populations to escape control efforts. To examine the population structure and the potential impacts of recent large-scale control interventions, we have investigated the genomic patterns of differentiation in mosquitoes belonging to the *Anopheles nili* group, a large taxonomic group that diverged ∼3-Myr ago. Using 4343 single nucleotide polymorphisms (SNPs), we detected strong population structure characterized by high *F*_ST_ values between multiple divergent populations adapted to different habitats within the Central African rainforest. Delineating the cryptic species within the *Anopheles nili* group is challenging due to incongruence between morphology, ribosomal DNA and SNP markers consistent with incomplete lineage sorting and/or interspecific gene flow. A very high proportion of loci are fixed (*F*_ST_ = 1) within the genome of putative species, which suggests that ecological and/or reproductive barriers are maintained by strong selection on a substantial number of genes.

## 1. Introduction

One of the principal goals of population genetics is to summarize the genetic similarities and differences between populations (Wright, 1984). This task can be relatively straightforward for some taxa, but the genetic relationship among populations can also be difficult to summarize, especially for species whose evolutionary history is complex and reticulate. The best known mosquito species of the genus *Anopheles* — which includes all vectors of *Plasmodium*, the parasite of human malaria — exhibit very complex range-wide population structure due to the combined effects of cryptic speciation, adaptive flexibility and ongoing gene flow (Harbach, 2013; Krzywinski and Besansky, 2003). For example, almost all major malaria vectors of the Afrotropical region belong to large taxonomic groups encompassing multiple incipient species relatively isolated reproductively and geographically from one another (reviewed by Sinka et al. 2010; Antonio-Nkondjio and Simard 2013; Coetzee and Koekemoer 2013; Dia et al. 2013; Lanzaro and Lee 2013). These characteristics make them promising model systems to study speciation and the processes which contribute to reproductive barriers (e.g. Turner et al. 2005; Lawniczak et al. 2010; Neafsey et al. 2010; Fontaine et al. 2015; Weng et al. 2016), but can also have far-reaching practical consequences. Both spatial and temporal variability in malaria cases and the effectiveness of vector control measures are greatly impacted by heterogeneity within vector species (Molineaux and Gramiccia 1980; Van Bortel et al. 2001). For these reasons, research on the genetic structure among the major African malaria vector mosquitoes has intensified over the last few decades (Antonio-Nkondjio and Simard, 2013; Coetzee and Koekemoer, 2013; Dia et al., 2013; Lanzaro and Lee, 2013).

The recent scaling up of insecticide-treated nets usage and indoor insecticide spraying to a lesser extent have led to a dramatic reduction in malaria morbidity across the continent (WHO, 2016). However, the other implications of these large-scale interventions include increased insecticide resistance (reviewed by Hemingway et al. 2016; Ranson and Lissenden 2016), range shift (e.g. Bøgh et al. 1998; Derua et al. 2012; Mwangangi et al. 2013) and profound evolutionary changes among vector populations. In contrast to insecticide resistance and range shift, which have been extensively studied, the recent adaptive changes among mosquito populations have yet to be addressed significantly. These changes — which involve local adaptation and genetic differentiation, introgressive hybridization and selective sweeps across loci conferring resistance to xenobiotics — are particularly evident in the most anthropophilic species (Barnes et al., 2017; Clarkson et al., 2014; Kamdem et al., 2017; Norris et al., 2015).

The ecology, taxonomic complexity, geographic distribution, role in transmission and evolutionary potential of each major African malaria vector are unique. Consequently, further research is needed to specifically resolve the fine-scale population structure and the genomic targets of natural selection in all of the important taxa including currently understudied species. The present work focused on a group of malaria vector species representing a large taxonomic unit named *Anopheles nili* group. Despite the significant role of some of its species in sustaining high malaria transmission, this group has received little attention. To date, four species that occur in forested areas of Central and West Africa and are distinguishable by slight morphological variations are known within the *An. nili* group: *An. nili sensu stricto* (hereafter *An. nili*), *An. ovengensis*, *An. carnevalei*, and *An. somalicus* (Awono-Ambene et al., 2004; Gillies and Coetzee, 1987; Gillies and De Meillon, 1968). These species are characterized by reticulate evolution and complex phylogenies that have been challenging to resolve so far (Awono-Ambene et al., 2004; Awono-Ambene et al., 2006; Kengne et al., 2003; Ndo et al., 2010, 2013; Peery et al., 2011; Sharakhova et al., 2013). Populations of *An. nili* and *An. ovengensis* are very anthropophilic and efficient vectors of *Plasmodium* in rural areas where malaria prevalence is particularly high (Antonio-Nkondjio et al., 2006).

To delineate genomic patterns of differentiation, we sampled mosquito populations throughout the range of species of the *An. nili* group in Cameroon and used reduced representation sequencing to develop genome-wide SNP markers that we genotyped in 145 individuals. We discovered previously unknown subpopulations characterized by high pairwise differentiation within *An. ovengensis* and *An. nili*. We further explored the genetic differentiation across the genome and revealed the presence of a very high number of outlier loci that are targets of selection among locally adapted subpopulations. These findings provide significant baseline data on the genetic underpinnings of adaptive divergence and pave the way for further genomic studies in this important group of mosquitoes. Notably, a complete reference genome will allow in-depth studies to decipher the functional and phenotypic characteristics of the numerous differentiated loci as well as the contribution of recent selective events in ongoing adaptation.

## 2. Materials and methods

### (a) Mosquito species

We surveyed 28 locations within the geographic ranges of species of the *An. nili* group previously described in Cameroon (Figure 1) (Antonio-Nkondjio et al., 2009; Awono-Ambene et al., 2004; Awono-Ambene et al., 2006; Ndo et al., 2010, 2013). The genetic structure of *Anopheles* species is most often based on macrogeographic or regional subdivisions of gene pools, but can also involve more subtle divergence between larvae and adults, or between adult populations found in or around human dwellings (e.g. Riehle et al., 2011). To effectively estimate the genetic diversity and identify potential cryptic populations within species, we collected larvae and adult mosquitoes within and around human dwellings using several sampling techniques (Service, 1993) in September-October 2013 (Table S1). To identify the four currently known members of the *An. nili.* group, we used morphological keys and a diagnostic PCR, which discriminates species on the basis of point mutations of the ribosomal DNA (Awono-Ambene et al., 2004; Gillies and Coetzee, 1987; Gillies and De Meillon, 1968; Kengne et al., 2003).

**Figure 1:**
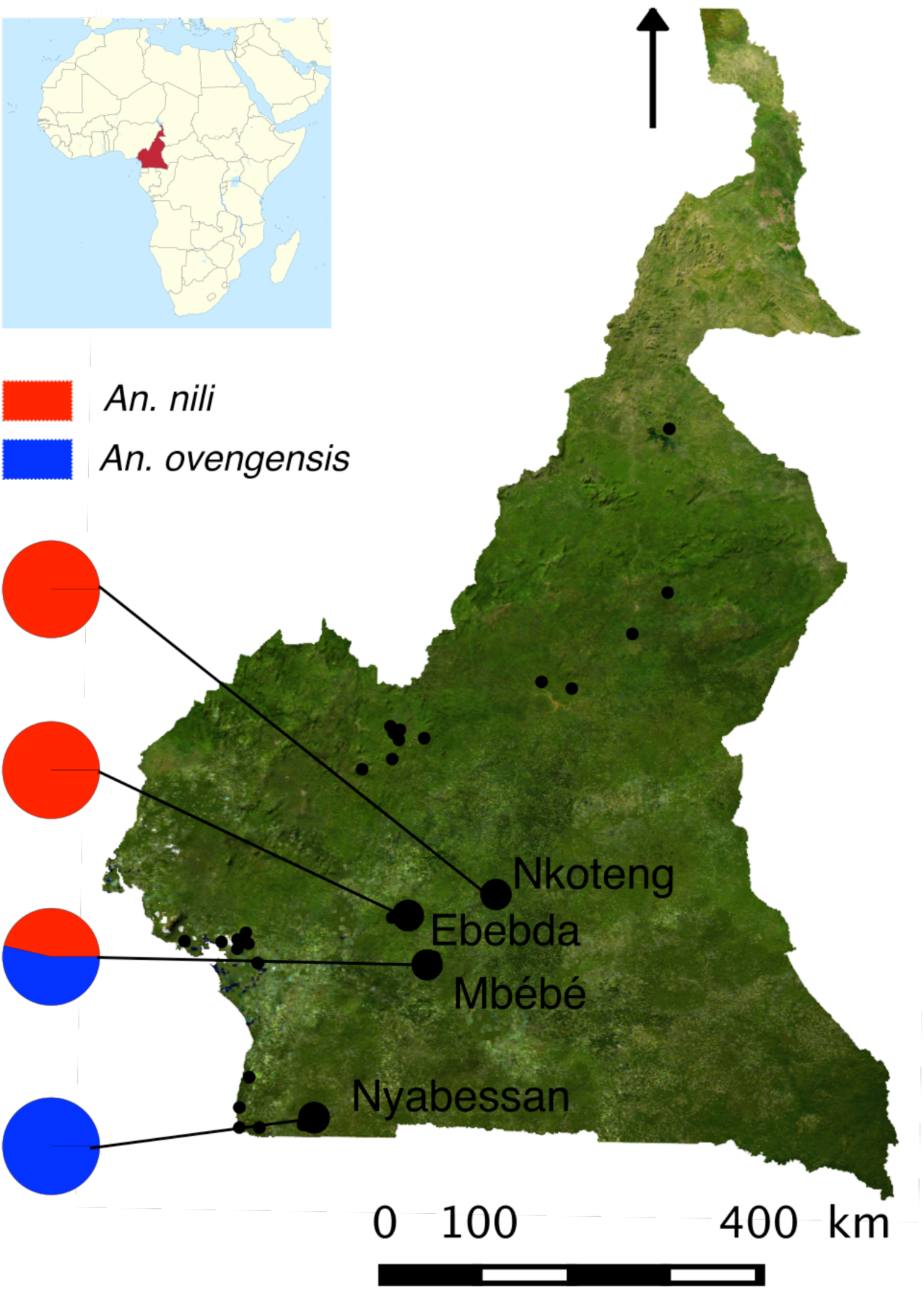
Map showing the sampling locations and relative frequencies of the morphological species *An. nili* and *An. ovengensis*. Small and large black dots indicate respectively the 28 locations surveyed and the four sampling sites where mosquitoes were collected.

### (b) Library preparation, sequencing and SNP discovery

We created double-digest Restriction site Associated DNA (ddRAD) libraries as described in Kamdem et al. 2017 using a modified version of the protocol designed by Peterson et al. 2012. Briefly, genomic DNA of mosquitoes was extracted using the DNeasy Blood and Tissue kit (Qiagen) and the Zymo Research MinPrep kit for larvae and adult samples respectively. Approximately 50ng (10μl) of DNA of each mosquito was digested simultaneously with *MluC1* and *NlaIII* restriction enzymes. Digested products were ligated to adapter and barcode sequences enabling identification of individuals. Samples were pooled, purified, and 400-bp fragments selected. The resulting libraries were amplified via PCR and purified, and fragment size distribution was checked using the BioAnalyzer. PCR products were quantified, diluted and single-end sequenced to 100 base reads on Illumina HiSeq2000.

### (b) SNP discovery and genotyping

The *process_radtags* program of the Stacks v 1.35 pipeline (Catchen et al., 2013; Catchen et al., 2011) was used to demultiplex and clean Illumina sequences. Reads that passed quality filters were aligned to the *An. nili* Dinderesso draft genome assembly (Giraldo-Calderon et al., 2015) made up of 51048 short contigs (∼200-30512bp long) using Gsnap (Wu and Nacu, 2010). To identity and call SNPs within consensus RAD loci, we utilized the *ref_map.pl* program of Stacks. We set the minimum number of reads required to form a stack to three and allowed two mismatches during catalogue creation. We generated SNP files in different formats for further downstream analyses using the *populations* program of Stacks and Plink v1.09 (Purcell et al., 2007).

### (d) Population genomics analyses

We analyzed the genetic structure of *An. nili* sensu lato (s.l.) populations using a Principal Component Analysis (PCA) and an unrooted Neighbor-Joining tree (NJ). We also examined ancestry proportions and admixtures between populations in Admixture v1.23 (Alexander et al., 2009) and Structure v2.3.4 (Pritchard et al., 2000). We used the package *adegenet* (Jombart, 2008) to implement the PCA in R (R Development Core Team 2016). The NJ tree was generated from SNP allele frequencies via a matrix of Euclidian distance using the R package *ape* (Paradis et al., 2004). We ran Admixture with 10-fold cross-validation for values of k from 1 through 8. Similarly, we analyzed patterns of ancestry from k ancestral populations in Structure, testing five replicates of k = 1-8. We used 200000 iterations and discarded the first 50000 iterations as burn-in for each Structure run. Clumpp v1.1.2 (Jakobsson and Rosenberg, 2007) was used to summarize assignment results across independent runs. To identify the optimal number of genetic clusters in our sample, we applied simultaneously the lowest cross-validation error in Admixture, the adhoc statistic deltaK (Earl and VonHoldt, 2012; Evanno et al., 2005) and the Discriminant Analysis of Principal Component (DAPC) method implemented in *adegenet.* To examine the level of genomic divergence among populations, we assessed genetic differentiation (*F*_ST_) across SNPs using the *populations* program of the Stacks pipeline. Mean *F*_ST_ values were also used to quantify pairwise divergence between populations. To infer the demographic history of different populations, we used the diffusion approximation method implemented in the package ∂a∂i v 1.6.3 (Gutenkunst et al., 2009). Single-population models were fitted to allele frequency spectra, and the best model was selected using the lowest likelihood and Akaike Information Criterion as well as visual inspections of residuals.

## Results

### (a) SNP genotyping

We collected mosquitoes from four locations out of 28 sampling sites (Figure 1, Table S1) and sequenced 145 individuals belonging, according to morphological criteria and diagnostic PCRs, to two species (*An. nili* (n = 24) and *An. ovengensis* (n = 121)). We assembled 197724 RAD loci that mapped to unique positions throughout the reference genome. After applying stringent filtering rules, we retained 408 loci present in all populations and in at least 50% of individuals in each population. Within these loci, we identified 4343 high-quality biallelic markers that were used to analyze population structure and genetic differentiation.

### (b) Morphologically defined species do not correspond to genetic clusters

PCA and a NJ tree show that the genetic variation across 4343 SNPs is best explained by more than two clusters, implying subdivisions within *An. nili* and *An. ovengensis* (Figure 2). Three subgroups are apparent within *An. nili* while two distinct clusters segregate in *An. ovengensis*. These five subpopulations are associated with the different sampling sites suggesting local adaptation of divergent populations. Importantly, Structure and Admixture analyses reveal that, at k = 2, one population identified by morphology and the diagnostic PCR as *An. nili* has almost the same ancestry pattern as the largest *An. ovengensis* cluster (Figure 3). Such discrepancies between morphology-based and molecular taxonomies can be due to a variety of processes including phenotypic plasticity, introgressive hybrydization or incomplete lineage sorting (i.e., when independent loci have different genealogies by chance) (Arnold, 1997; Combosch and Vollmer, 2015; Fontaine et al., 2015; Weng et al., 2016). At k = 2 and k = 3, some populations also exhibit half ancestry from each morphological species suggestive of gene flow. We found a conflicting number of genetic clusters in our samples likely reflecting the complex history of subdivisions and admixtures among populations (Figure 4). The Evanno et al., (2005) method, which highlights the early states of divergence between *An. nili* and *An. ovengensis* indicates two probable ancestors. DAPC and the Admixture cross-validation error, which are more sensitive to recent hierarchical population subdivisions, show five or more distinct clusters as revealed by PCA and the NJ tree (Figure 4).

**Figure 2:**
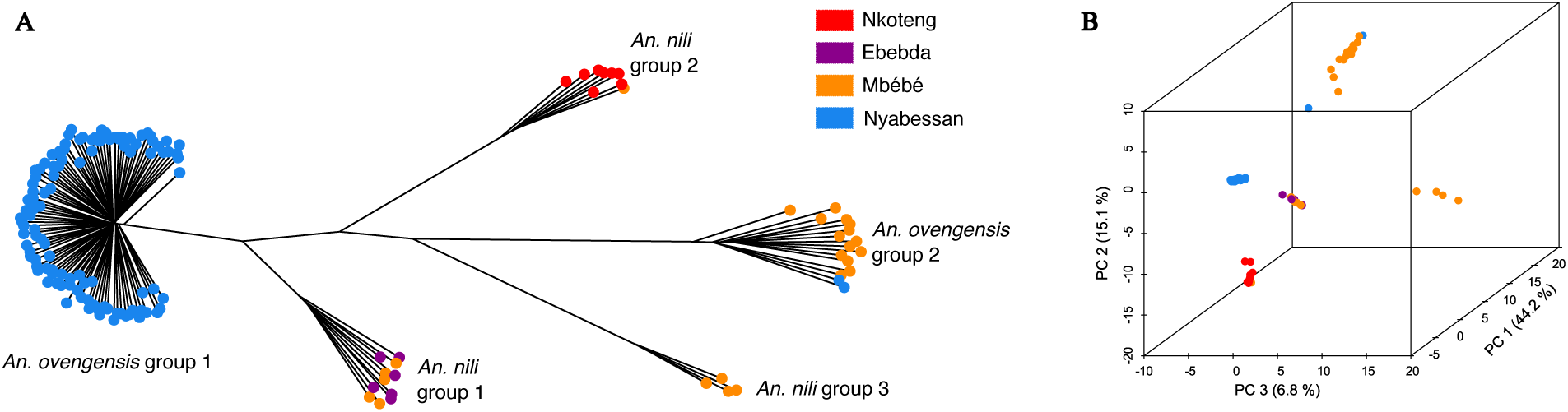
Population genetic structure inferred from 4343 SNPs using PCA (A) and a neighbor-joining tree (B). The percentage of variance explained is indicated on each PCA axis.

**Figure 3:**
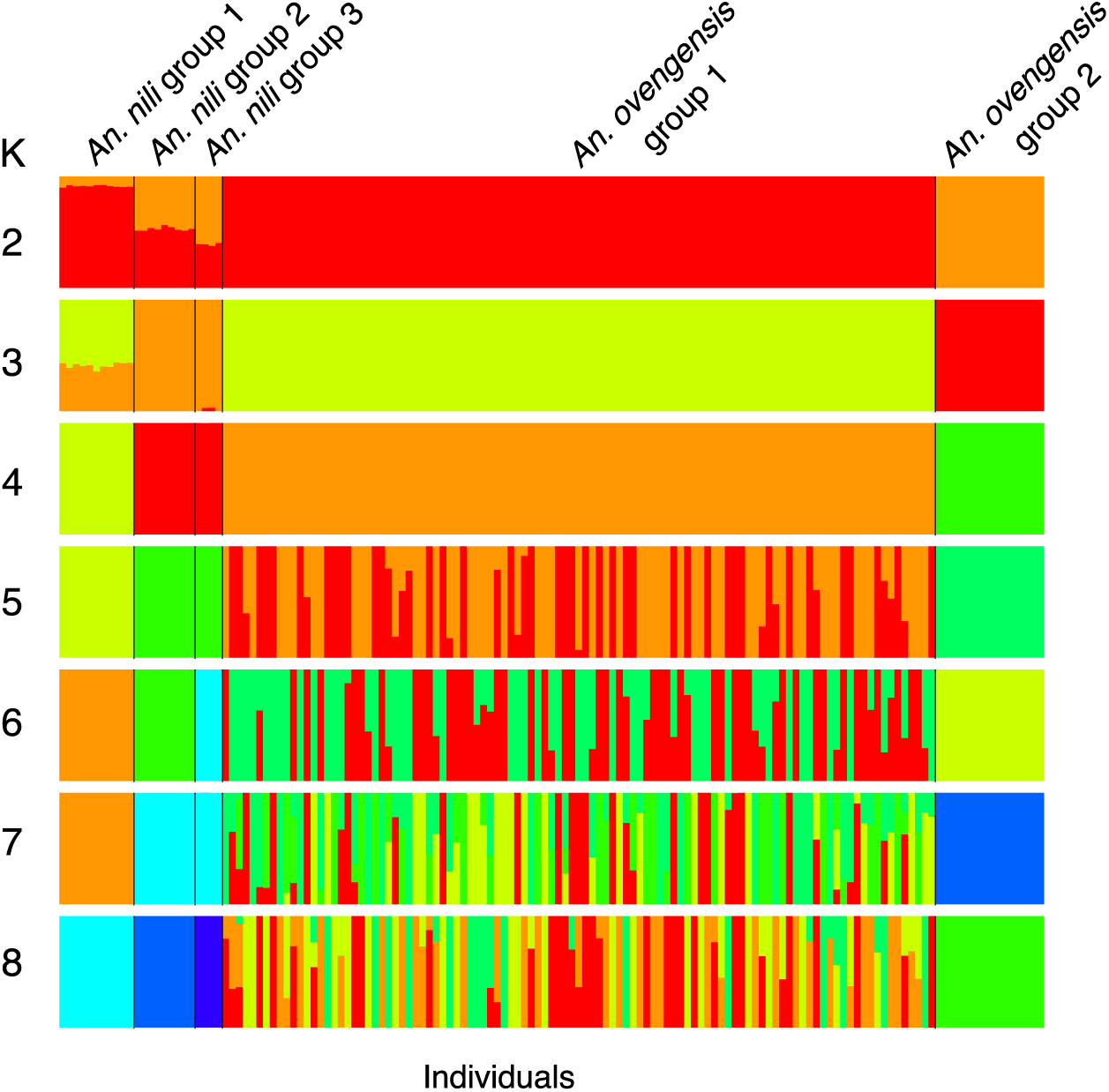
Ancestry proportions inferred in Admixture with k = 2 – 8.

**Figure 4:**
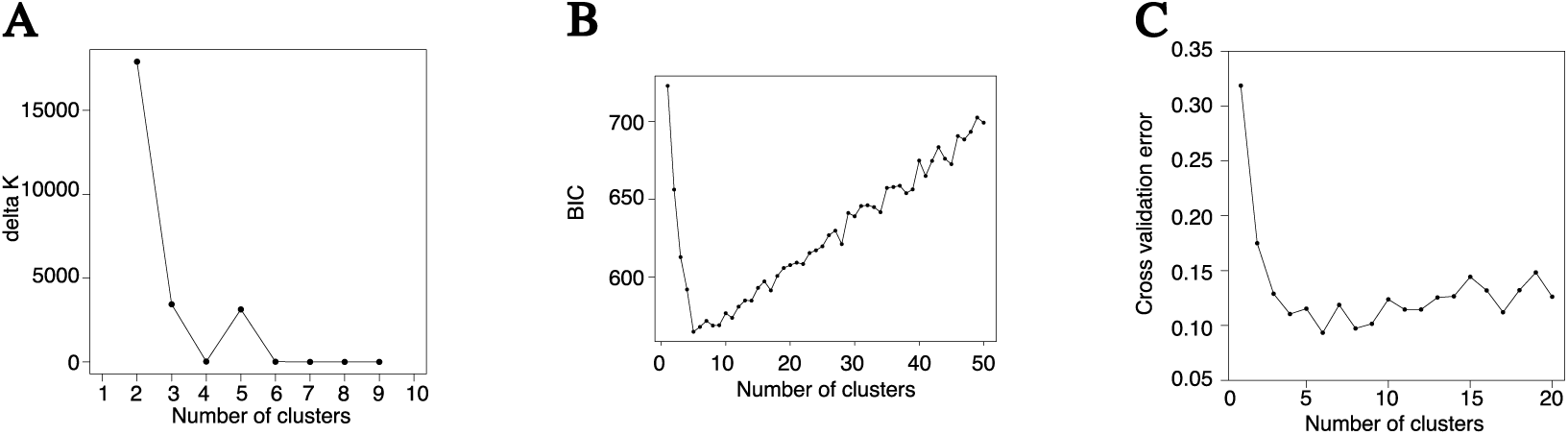
Identification of the optimal number of genetic clusters using the delta k method of Evanno et al., (2005) (A), DAPC (B) and 10-fold cross-validation in Admixture (C). The lowest Bayesian Information Criterion (BIC) and cross-validation error and the highest delta k indicate the most probable number of clusters.

As suggested by the long internal branches, which connect subpopulations on the NJ tree, there is strong differentiation between and within morphological species characterized by globally high *F*_ST_ values (Table 1). Relatively lower *F*_ST_ values observed between certain clusters may be due to greater inter-population migration and intermixing or more recent divergence. The *F*_ST_ values do not reflect the morphological delimitation of species, and the level of genetic differentiation is even higher between some subpopulations within the same morphological species. Overall, patterns of genetic structure and differentiation reveal a group of populations whose phylogenies and species status are likely confounded by hybridization and/or incomplete lineage sorting. We argue that the current taxonomy based on morphology and ribosomal DNA does not capture the optimal reproductive units among populations of this group of mosquitoes.

**Table 1:** Pairwise *F*_ST_ between divergent subpopulations of *An. nili* s.l.

| $F_{ST}$ | *An. nili*<br>group 1 | *An. nili*<br>group 2 | *An. nili*<br>group 3 | *An.*<br>*ovengensis*<br>group 1 | *An.*<br>*ovengensis*<br>group 2 |
| --- | --- | --- | --- | --- | --- |
| *An. nili* group 1 | - |  |  |  |  |
| *An. nili* group 2 | 0.374 | - |  |  |  |
| *An. nili* group 3 | 0.506 | 0.552 | - |  |  |
| *An. ovengensis*<br>group 1 | 0.135 | 0.275 | 0.364 | - |  |
| *An. ovengensis*<br>group 2 | 0.432 | 0.458 | 0.492 | 0.349 | - |

### (c) Genomic signatures of divergent selection and demographic history

We analyzed patterns of genetic differentiation across SNP loci throughout the genome. Pairwise comparisons are based on filtered variants that satisfy all criteria to be present in both populations, which explains the discrepancy in the number of SNPs observed between specific paired comparisons (Figure 5). The distribution of locus-specific *F*_ST_ values between the five subpopulations revealed a U-shape characterized by two peaks around 0 and 1. The great majority of SNPs have low to moderate divergence, but a substantial number of variants are extremely differentiated between populations. The maximum *F*_ST_ among SNPs is 1 and the proportion of loci with *F*_ST_ = 1 varies from 6.52% between the populations we termed *An. nili* group 1 and *An. ovengensis* group 1 to 44.74% between the subgroups called *An. nili* group 2 and *An. nili* group 3 (Figure 5). This pattern of genome-wide divergence suggests that a very high number of sites with abrupt differentiation — which likely contain genes that contribute to divergent selection and/or reproductive isolation — coexist with regions of weak divergence that can be freely exchanged between species. As is the case with the overall genetic differentiation, morphology is not a reliable predictor of locus-specific divergence. Precisely, the lowest percentage of fixed SNPs is found between *An. ovengensis* from Nyabessan and *An. nili* collected from Mbébé and Ebebda (Figure 1, Figure 5). In contrast, the greatest proportion of fixed loci is observed between locally adapted subgroups within the same morphological species, *An. nili*. The draft reference genome made up of short contigs did not allow us to test hypotheses about the genomic distribution of differentiated loci. For example, it remains unknown whether the numerous SNPs that are fixed among populations are spread throughout the entire genome or clustered in genomic regions of low recombination including chromosomal inversions and chromosome centers (Nosil and Feder, 2012; Roesti et al., 2012).

**Figure 5:**
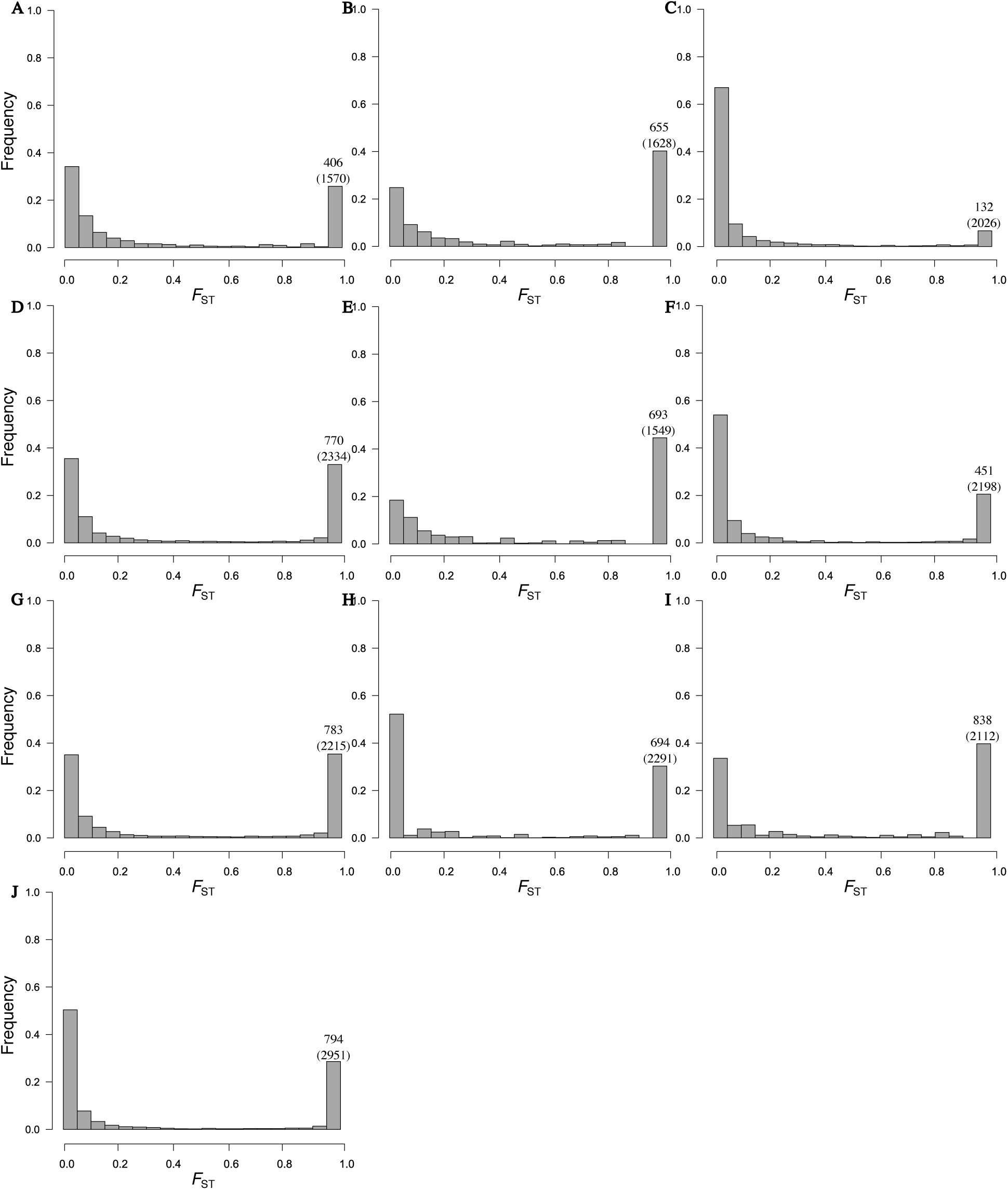
Distribution of *F*_ST_ values throughout the genome between *An. nili* group 1 and *An. nili* group 2 (A); *An. nili* group 1 and *An. nili* group 3 (B); *An. nili* group 1 and *An. ovengensis* group 1 (C); *An. nili* group 1 and *An. ovengensis* group 2 (D); *An. nili* group 2 and *An. nili* group 3 (E); *An. nili* group 2 and *An. ovengensis* group 1 (F); *An. nili* group 2 and *An. ovengensis* group 2 (G); *An. nili* group 3 and *An. ovengensis* group 1 (H); *An. nili* group 3 and *An. ovengensis* group 2 (I); *An. ovengensis* group 1 and *An. ovengensis* group 2 (J). The number of SNPs with *F*_ST_ = 1 is indicated in each pairwise comparison as well as the total number of SNPs in parenthesis.

Models of population demography indicate that all subgroups have experienced an increase in effective size in a more or less recent past (Table 2). Nevertheless, confidence intervals of population parameters are high in some populations, and our results should be interpreted with the necessary precautions. The population growth is less significant in *An. nili* group 1.

**Table 2:** Demographic models of different subgroups of *An. nili* s.l.

| Population | Best Model | Log Likelihood | Final Population<br>Size<sup>a</sup> (95% CI) | Time<sup>b</sup> (95% CI) |
| --- | --- | --- | --- | --- |
| *An. nili* group 1 | *Growth* | -18.42 | 6.41 (5.326 - 20.71) | 3.70 (1.11 - 13.31) |
| *An. nili* group 2 | *Two-epoch* | -19.97 | 17.87 (9.33 - 35.50) | 11.27 (4.93 - 19.64) |
| *An. ovengensis* group 1 | *Growth* | -112.18 | 13.04 (12.15 - 17.26) | 0.70 (0.58 - 1.08) |
| *An. Ovengensis* group 2 | *Growth* | -22.98 | 19.95 (14.45 - 45.70) | 5.11 (2.33 - 15.13) |

<sup>a</sup> Relative to ancestral population size.
<sup>b</sup> Expressed in units 2Ne generations from start of growth to present.

## Discussion

### (a) Genetic differentiation

Advances in sequencing and analytical approaches have opened new avenues for the study of genomes of disease vectors. We have focused on malaria mosquitoes of the *An. nili* group, whose taxonomy and population structure have been challenging to resolve with low-resolution markers. We analysed the genetic structure using genome-wide SNPs and found strong differentiation and local adaption among populations belonging to the two morphologically defined species *An. nili* and *An. ovengensis*. The exact number of subpopulations remains contentious, with the suggested number of divergent clusters varying from two to five within both species. Significant population structure at eight microsatellite loci has been described among *An. nili* populations from Cameroon, with *F*_ST_ values as high as 0.48 between samples from the rainforest area (Ndo et al., 2013). By contrast, *An. ovengensis* was discovered recently and the genetic structure of this vector remains understudied. This species was initially considered as a sibling of *An. nili* (Awono-Ambene et al., 2004; Awono-Ambene et al., 2006; Kengne et al., 2003), but more recent studies have started to challenge the assumed relatedness between the two species due to the high divergence of their polytene chromosomes (Sharakhova et al., 2013). Our findings call for careful review of the current taxonomy within this group of species, which is a necessary first step for accurately delineating the role played by the different subpopulations in malaria transmission.

Our samples were collected from locations characterized by a more or less degraded forest in the rainforest area of Cameroon. In these habitats, larvae of *An. nili* s.l. exploit relatively similar breeding sites consisting of slow-moving rivers (Antonio-Nkondjio et al., 2009). The ecological drivers of genetic differentiation remain unknown, and will be difficult to infer from our data given the apparent similarity of habitats among populations. Further study is needed to clearly address the environmental variables that may be correlated with ongoing adaptive divergence at adult and larval stages. One of the most expected outcomes of current large-scale malaria control measures that are underway in Sub-Saharan African countries concerns the effects of increased insecticide exposure on the genetic diversity and population demography of vectors. A substantial population decline that may considerably affect the adaptive potential of vector species has been occasionally reported following the distribution of insecticide-treated bed nets and/or indoor residual house spraying (e.g. Athrey et al., 2012). The inferred demographic history of the different subpopulations within the *An. nili* group does not reveal signatures of bottlenecks that can be potentially correlated with increased usage of insecticides and insecticide-treated nets. This result is consistent with the demography of several other important malaria vectors of the Afrotropical region, including *An. gambiae*, *An. coluzzii*, *An. funestus* and *An. moucheti*, which reveals a substantial population increase suggesting that recent intense insecticide exposure has yet to leave deep or detectable impacts on patterns of genetic variation among mosquito populations (Fouet et al., 2017; Kamdem et al., 2017; O’Loughlin et al., 2014).

### (b) Genomic architecture of geographic and reproductive isolation

Understanding the genomic architecture of reproductive isolation may reveal crucial information on the sequence of events that occur between the initial stages of divergence among populations and the onset of strong reproductive barriers (e.g. Turner et al., 2005; Harr 2006; Nadeau et al., 2012; Ellegren et al., 2012; Carneiro et al., 2014; Burri et al., 2015). One influential concept of speciation coined the “genic view of species” proposes that boundaries between species are properties of individual genes or genome regions and not of whole organisms or lineages (Barton and Hewitt, 1985; Harrison and Larson, 2014; Harrison, 1990; Key, 1968; Nosil and Feder, 2012; Rieseberg et al., 1999; Wu, 2001). We have discovered a dramatically high number of SNPs that are strongly differentiated between populations and often fixed within subgroups of *An. nili* s.l. Interpreting this intriguing pattern of genomic differentiation is not straightforward due to the complex interactions between numerous forces — including positive or negative selection, recombination, introgressive hybridization and incomplete lineage sorting — that can affect the level of divergence among SNPs (Begun and Aquadro, 1992; Cutter and Payseur, 2013; Harrison and Larson, 2016; Nachman and Payseur, 2012; Roesti et al., 2012). Some of these variants exhibiting high divergence among populations certainly contain markers of ecological and/or reproductive isolation. However, as far as reproductive barriers are concerned, recent studies have indicated a complex relationship between genetic differentiation and gene flow at the genome level (e.g. Gompert et al., 2012; Hamilton et al., 2013a, b; Larson et al., 2013, 2014; Parchman et al., 2013; Taylor et al., 2014). Highly divergent genomic regions do not necessarily coincide with regions of reduced gene flow. Several alternative interpretations exist for the numerous high-*F*_ST_ regions we detected in all pairwise comparisons (Cruickshank and Hahn, 2014; Delmore et al., 2015; Nachman and Payseur, 2012; Noor and Bennett, 2009). Nevertheless, careful examination of these outliers of differentiation may reveal significant insights into the wide range of genes and traits that contribute to ecological divergence and/or reproductive isolation between subgroups of *An. nili* s.l. A complete genome assembly will be necessary to better delineate specific regions of the genome under natural selection, and thereby clarify the genomic basis of phenotypic fitness differences between divergent populations. This will also help understand the extent to which recent selection associated with human interventions contribute to local adaptation and genetic differentiation as observed in *An. gambiae* and *An. coluzzii* (Kamdem et al., 2017).

Signals consistent with gene flow between *An. nili* and *An. ovengensis* are apparent in our data despite significant time since divergence (∼3M-yr) (Ndo et al., 2013). Some individuals display almost half ancestry from each morphological species. The disagreement between morphology/PCR and molecular taxonomies observed in Structure and Admixture analyses also suggests that incongruent genealogies may be widespread along chromosomes due to hybridization. However, hybridization can be difficult to detect because other factors such as incomplete lineage sorting or technical artefacts can leave signatures that are similar to those of interspecific gene flow (Liu et al., 2014; Patterson et al., 2012). A complete reference genome is also needed to analyze the detailed distribution of genealogies across small genomic windows and to disentangle the relative contribution of processes that generate the putative admixtures and species confusion observed among divergent populations (Fontaine et al., 2015; Martin et al., 2013; Weng et al., 2016).

## 5. Conclusions and implications

Delineating the fine-scale population structure of mosquito populations is crucial for understanding their epidemiological significance and their potential response to vector control measures. Moreover, recent malaria control efforts are affecting interspecific gene flow, genetic differentiation, population demography and natural selection in mosquitoes (Athrey et al., 2012; Barnes et al., 2017; Clarkson et al., 2014; Kamdem et al., 2017; Norris et al., 2015). Deciphering the signatures of these processes across mosquito genomes is important to minimize their negative impacts on vector control. Our findings shed some light on the complex evolutionary history and provide a framework for future investigations into the genetic basis of ecological and reproductive barriers among species of the *An. nili* group.

## Acknowledgements

Funding for this project was provided by the University of California Riverside and NIH grants 1R01AI113248 and 1R21AI115271 to BJW. We thank inhabitants and administrative authorities of the sampling sites included in this study for their collaboration.

### Author contributions

Conceived and designed the experiments: CF CK BJW. Performed the experiments: CF CK SG BJW. Analysed the data: CF CK. Wrote the paper: CF CK BJW.

### Data Archiving Statement

Data for this study are available at: to be completed after manuscript is accepted for publication

## Supplemental information

**Table S1:** Information on mosquito samples included in this study.

| Sampling locations | Geographic coordinates | Sampling methods |  |  | Total |
| --- | --- | --- | --- | --- | --- |
|  |  | HLC-OUT | HLC-IN | LC |  |
| Ebebda | 4°20'00"N, 11°17'00"E |  |  | 6 | 6 |
| Nkoteng | 4°31'00"N, 12°02'00"E |  |  | 8 | 8 |
| Nyabessan | 2°24'00"N, 10°24'00"E | 63 | 44 |  | 107 |
| Mbébé | 4°10'00"N, 11°04'00"E | 13 | 3 | 8 | 24 |
| Total |  | 76 | 47 | 22 | 145 |

HLC-OUT, human landing catches performed outdoor; HLC-IN, human landing catches performed indoor; LC, larval collection

